# Proteomic Dynamics and Heat Stress Response in *Arabidopsis* Seedlings

**DOI:** 10.1101/2024.04.30.591943

**Authors:** Kai-Ting Fan, Yuan Xu

## Abstract

Global warming poses a grave threat to plant survival, adversely affecting growth and agricultural productivity. To develop thermotolerant crops, a profound comprehension of plant responses to heat stress at the molecular level is imperative. Leveraging a novel fusion of 15N-stable isotope labeling and the ProteinTurnover algorithm, we meticulously investigated proteome dynamics in Arabidopsis thaliana seedlings subjected to moderate heat stress (30°C). This innovative approach facilitated a comprehensive analysis of proteomic changes across diverse cellular fractions. Our study unveiled significant turnover rate alterations in 571 proteins, with a median increase of 1.4-fold, indicative of accelerated protein dynamics under heat stress. Notably, root soluble proteins exhibited more subdued changes, suggesting tissue-specific adaptations. Moreover, we observed noteworthy turnover variations in proteins associated with redox signaling, stress response, and metabolism, underscoring the complexity of the response network. Conversely, proteins involved in carbohydrate metabolism and mitochondrial ATP synthesis displayed minimal turnover changes, signifying their stability. This exhaustive examination sheds light on the proteomic adjustments of Arabidopsis seedlings to moderate heat stress, elucidating the delicate balance between proteome stability and adaptability. These findings significantly augment our understanding of plant thermal resilience and offer crucial insights for the development of crops endowed with enhanced thermotolerance.

## 1. Introduction

High temperature is one of the most deleterious abiotic stresses for plants as it affects many aspects of plant growth, reproduction, and yield. Greenhouse gases have been greatly elevated since the Industrial Revolution resulting in global warming, and there is a greater than 90% chance that by the end of the 21st century, the average growing season temperatures in the tropics and subtropics will exceed the highest temperature on record (1990-2006) [1]. With this warming climate, the development of crop cultivars engineered for improved thermotolerance [2,3] is needed to insure the food supply. At the whole plant level, heat stress induces observable phenotypes such as suppressed seed germination, inhibited shoot and root growth, fruit discoloration, leaf senescence, and reduced yield [4]. At the cellular level, heat stress leads to physical perturbations like increased membrane fluidityand protein denaturation, affectingprotein synthesis, enzyme activity, and metabolism [5,6]. Conversely, moderate heat stress, such as 28°C, induces phenotypes suggesting enhanced evaporative cooling capacity, despite increased water loss and transpiration rates [7].

Photosynthesis, particularly Photosystem II (PSII), is significantly affected by heat stress, with moderate heat causing PSII photoinhibition [8] and higher temperatures leading to dissociation or inhibition of the oxygen-evolving complex [9]. While Rubisco, the enzyme responsible for carbon fixation, is inherently thermostable in higher plants, heat stress can inhibit Rubisco activase, thereby impacting carbon assimilation rates [10,11]. Research by Kurek et al. highlights Rubisco activase as a major limiting factor in plant photosynthesis under heat stress, with the introduction of thermostable Rubisco activase variants resulting in increased carbon assimilation rates under moderate high temperatures [10].

Plants employ multiple molecular mechanisms to adapt to elevated ambient temperatures. Elevated temperatures increase the concentration of misfolded, unfolded, and aggregated proteins, leading to the transcriptional activation of heat stress-induced genes [12]. These genes include various families of heat shock proteins (HSPs), which function as molecular chaperones controlling protein folding and stability [2]. The unfolded protein response (UPR) in plants is a vital signaling pathway in response to stress, triggering processes including protein translation attenuation, activation of the ER-associated degradation pathway, and induction of endoplasmic reticulum (ER) chaperones [13]. As heat stress affects protein stability, it also disrupts specific enzyme functions, perturbing metabolism. Oxidative stress accompanies the heat stress response, leading to the accumulation of reactive oxygen species (ROS). Coping with the accumulation of ROS and other oxidative stress injuries is a major challenge for organisms facing heat stress. ROS production triggers an antioxidant response mediated through a MAPK signal pathway and induction of downstream transcription factors. A key aspect of this response involves removing ROS molecules using ROS scavenging enzymes such as ascorbate peroxidase (APX) and catalase (CAT) [12].

In the field of genomics, researchers have identified thousands of genes that may be differentially regulated at the transcriptional level in response to heat stress in various plant species, including *Arabidopsis* [14], tomato [15], rice [16], barley [17], wheat [18], and maize [19]. However, the steady-state levels of transcripts do not fully reflect the levels of corresponding proteins, as translation serves as a crucial point of regulatory control in the plant heat stress response [20,21]. These studies underscore the inadequacy of solely relying on transcriptional analyses of the heat response in plants.

With continuous advancements in liquid chromatography (LC) coupled mass spectrometry (MS) instrumentation over the past two decades, proteomics has enabled new approaches for analyzing protein abundance and dynamics in response to stress conditions [22]. However, based on our knowledge, relatively few have explored the effects of stress conditions on protein dynamics or turnover. Isotopic labeling techniques have become indispensable tools for investigating turnover dynamics within plant systems[23,24]. One such example is documented by Li et al. [25], who utilized ^15^N-labeling and two-dimensional fluorescence difference gel electrophoresis with LC-MS/MS to measure the protein degradation rates of 84 proteins in *Arabidopsis* suspension cells. They then calculated the protein synthesis rate based on degradation rates and changes in protein relative abundance. The study concluded that protein turnover rates generally correlated with protein function and among protein complex subunits. Proteins associated with RNA/DNA binding and metabolism, protein synthesis and degradation, and stress and signaling exhibited higher degradation and synthesis rates, while those associated with antioxidant and defense mechanisms, mitochondrial energy metabolism, and primary metabolism had lower rates. Within these functional categories, the stress and signaling category displayed the highest average degradation and synthesis rate. Furthermore, the relative degradation and synthesis rates were examined to determine which proteins would experience changes in abundance due to alterations in their turnover dynamics. The study found a positive correlation between synthesis and degradation rates for proteins involved in antioxidant defense and protein synthesis and degradation categories, but no correlation for mitochondrial energy, primary metabolism, or stress and signaling proteins. This suggests a tendency to maintain stable levels of proteins in the antioxidant defense and protein synthesis and degradation categories while allowing for rapid responses of cytosolic and nuclear proteins to environmental changes or stress. Specific proteins, such as glutathione peroxidase 6 involved in antioxidant stress defense, heat-shock protein 60 (HSP60) involved in protein folding, and the glutathione S-transferase Phi family involved in detoxification, exhibited slow degradation rates. Moreover, mitochondrial proteins were generally more stable than cytosolic and nuclear proteins, indicatinga preference for maintaining stable mitochondrial protein function while allowing for the dynamic adjustment of cytosolic and nuclear proteins to environmental stimuli or stressors.

Using a similar approach, Nelson et al. further measured the degradation rate (Kd, day-1) of 224 mitochondrial proteins using 7-day-old *Arabidopsis* cell culture with 1, 4, 5, and 7 days of ^15^N-label incorporation [26]. Both studies utilized the Isodist algorithm [27] to assign the isotopic abundance of natural abundance and labeled peptide mass spectral data to obtain Relative Isotope Abundance (RIA) values for each peptide throughout the time course. However, for each replicate, a protein’s RIA at a given time point was calculated as the median of all measured peptide RIA values for the corresponding protein. The average RIA value across all replicates was then used as the given protein’s RIA at each time point. The protein degradation rate was computed from the slope coefficient of the linear regression of the natural logarithm of RIA against time. Although this method is convenient for estimating proteome degradation rates in rapidly growing cellular systems, the higher complexity of multicellular or slow-growing organisms, coupled with the difficulties in interpreting overlapping isotopic distributions in partially labeled systems, limits the applicability of the approach to intact organisms. Another disadvantage is that, in order to detect significant changes in protein turnover rates across different conditions or treatments, the individual contributions of specific peptides to the overall protein turnover are lost due to the use of median peptide RIA values for each protein. This unnecessarily discards potentially important information regarding the inherent heterogeneity of intracellular protein populations.

Here, a proteome-wide analysis was conducted to monitor changes in proteome turnover of *Arabidopsis thaliana* seedling tissues after exposure to elevated temperature (30°C). This study presents a novel approach to evaluate, for the first time, the contribution of the dynamic balance of protein synthesis and degradation in response to moderate heat stress in intact plant seedlings. The algorithm *ProteinTurnover* were used to measures protein turnover rates using ^15^N-metabolic stable isotope labeling approach on a proteomic scale [28]. In this study, hundreds of proteins have been identified in root or shoot soluble, organellar, and microsomal fractions with significant changes in turnover rates in response to elevated temperature stress.

## 2. Results

The goal of this study was to assess how moderate heat treatment influences protein turnover rates across various cellular fractions of *Arabidopsis* seedling tissues on a proteomic scale. To achieve this objective, the seedlings transferred to medium containing stable isotope ^15^N were grown under heat stress (30°C), whereas seedlings under control temperature (22°C) were continuously grown on ^14^N-medium. Root and shoot tissues harvested at 5 time points (0, 8, 24, 32, and 48 hours) post-transfer and subjected to differential centrifugation to isolate fractions enriched in organellar, soluble, or microsome-associated proteins were analyzed by LC-MS/MS.

### 2.1. Peptide Identification and Selection Criteria for Protein Turnover Rate Measurements

From the root tissue, 822 and 857 proteins were identified in the enriched soluble fraction from the control and 30°C groups, respectively. In the enriched organelle fraction, 494 and 377 proteins were identified from the control and 30°C groups, respectively. Additionally, 1,222 and 1,054 proteins were identified in the enriched microsomal fraction from the control and 30°C groups, respectively. At the time of analyzing these samples, a nano-LC inlet was not available. To compensate for this limitation, larger total quantities of protein were isolated, processed, and analyzed using a 2.1 mm UHPLC column and flow rates of 300 µL/min. Thousands of identified peptides were required for the subsequent turnover analysis due to the lower sensitivity inlet used in this study. As indicated in Table S-1, each sample contained between 5,000 to 14,000 peptides, but only 30 −50% of them were present in a sufficient number of time points to compute turnover rates. In this dataset, peptides were most frequently excluded because they were not identified in the time 0 dataset.

Applying multiple quantitative quality criteria for the inclusion of each peptide can enhance the qualityof the resultingturnover data and accelerate data processing. Peptides with significant standard errors typically represent those with poor spectral fitting, often due to co-eluting contaminants (Figure 1, panel A). Peptides were included in further analysis if they met specific criteria: a visual score for spectral fitting (to the beta-binomial model) greater than 80 out of 100, a standard error in the turnover rate fitting of less than 10, and data points for at least 3 of the time points (including time 0). These criteria were chosen based on empirical visual inspection of peptide turnover fitting plots generated by the algorithm. Additionally, the normal quantile-quantile (Q-Q) plot of peptide log_2_*k* was utilized to assess whether the log_2_*k* data were normally distributed (Figure 1, panels B and D). Scatter plots of log_2_*k* and the standard error of log_2_*k* (such as shown in Figure 1, panels A and C) aided in assessing dataset quality. Inspection of Figure 1 panel C also suggests a potential negative linear correlation between log_2_*k* and the standard error of log_2_*k*, at least for this dataset. Nonetheless, only peptides selected using the aforementioned filtering criteria were used for further turnover rate analysis. Once a peptide passed this filter, it was assumed that the turnover rate calculated for each peptide contributed equally to the final protein turnover rate. Therefore, the log_2_*k* of all selected peptides was averaged to yield each individual protein turnover rate (log_2_*k*) for a given experimental condition (control vs. treatment).

**Figure 1.**
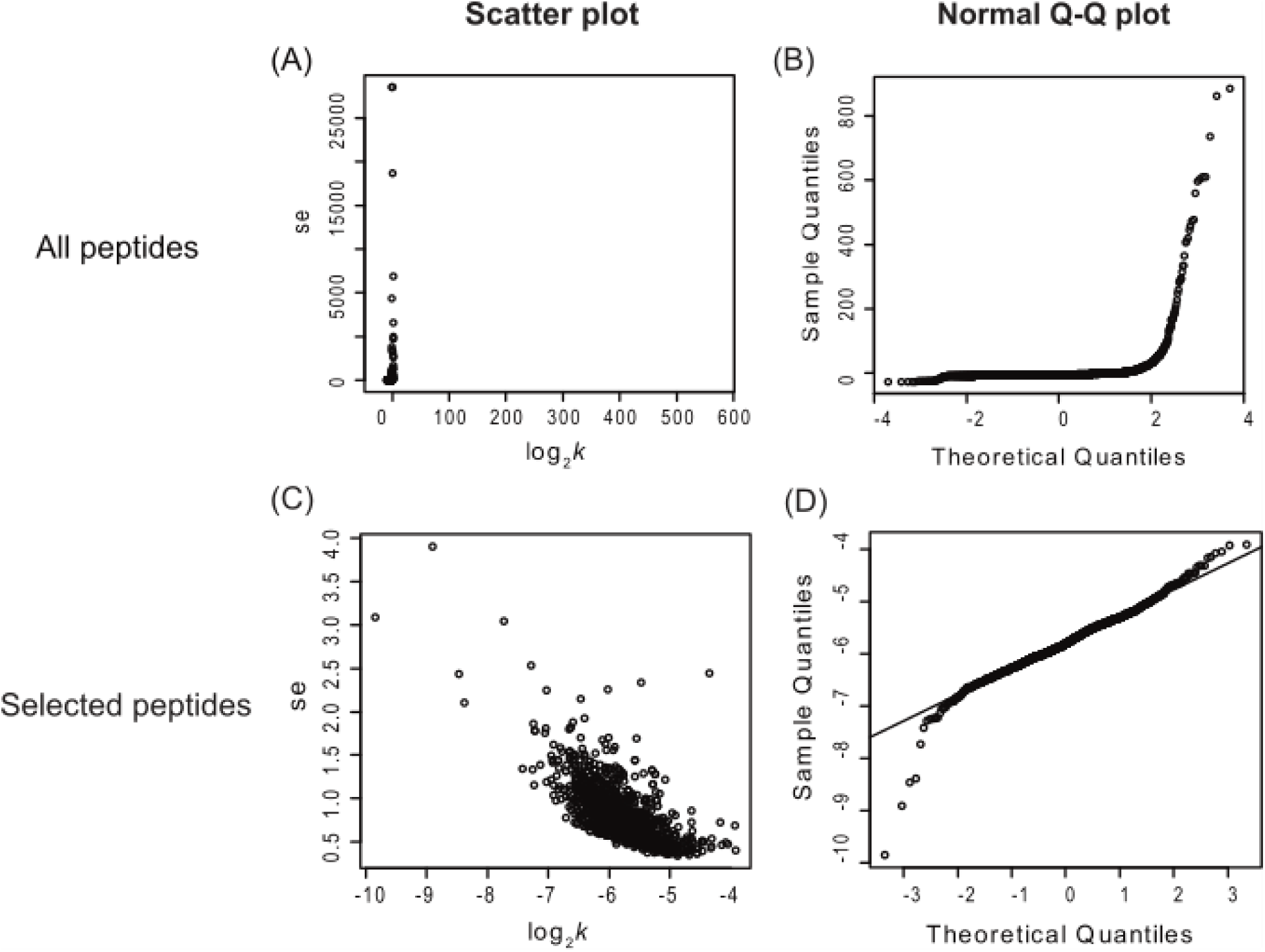
Scatter plots and Normal Q-Q plots of all identified *Arabidopsis* peptides (top) vs. peptides selected with visual scores higher than 80, standard error lower than 10, and at least 3 labeling time points (bottom). The panels on the left (A and C) are scatter plots of standard error of log_2_*k* (se.log_2_*k*) against log_2_*k*; the panels on the right (B and D) are normal Q-Q plots of each peptide’s turnover rate (log_2_*k* values). This figure shows only the peptide data from the enriched shoot soluble fraction and includes data combined from both control and heat treatment groups. The number of peptide n=10,400 (A and B) and 1,273 (C and D). ‘se’ = standard error.

### 2.2. Overview of the Effects of Heat Stress on Peptide and Protein Turnover Rates

#### 2.2.1. Trends of peptide or protein turnover rates

The distributions of peptide turnover rates (log_2_*k*) between the control and 30°C groups are depicted for comparison purposes as histograms for soluble, organellar, and microsomal protein-enriched fractions of shoot and root tissues in Figure 2. The distributions of protein turnover rates (log_2_*k*) between the control and 30°C groups are illustrated for comparison purposes as histograms for soluble, organellar, and microsomal protein-enriched fractions of shoot and root tissues in Figure 3. When comparing the mean values of peptide turnover rates or the median value of protein turnover rates between roots and shoots, generally across all fractions, the turnover rates of roots were faster than those of shoots. The average protein turnover rate (log_2_*k*) was −5.308, −5.594, and −5.377 in the soluble, organellar, and microsomal fractions, respectively, while in shoots, the average protein turnover rate was −6.0348, −6.1046, and −5.9765 in the soluble, organellar, and microsomal fractions, respectively. For the control group, the mean protein turnover rates (log_2_*k*) were close to −5.39 in roots and −6.03 in shoots, indicating that the mean protein half-lives were 29.13 hours in roots and 45.2 hours in shoots, suggesting that root proteome might have a faster turnover rate than shoot proteome in general. This may be related to the development of root tissue in the young seedling stage of plants, which requires more rapid changes in protein synthesis and degradation. For example, the fast turnover rate of plasma membrane proton pump (ATPase 1) (Table 1) suggests that the establishment of protein machinery for metabolite uptake could be essential for growth at this stage. Although several proteins had dramatically long half-lives (Table 1), the average protein turnover rates measured in this study were much faster than the average protein turnover rates in 21 to 26-day-old adult *Arabidopsis* leaves (∼4.6 days) as reported in the unpublished work from Millar et al. (presented at the 2015 ASPB conference), suggesting that more rapid protein turnover may be required in the seedling than the adult stage in plants.

**Figure 2.**
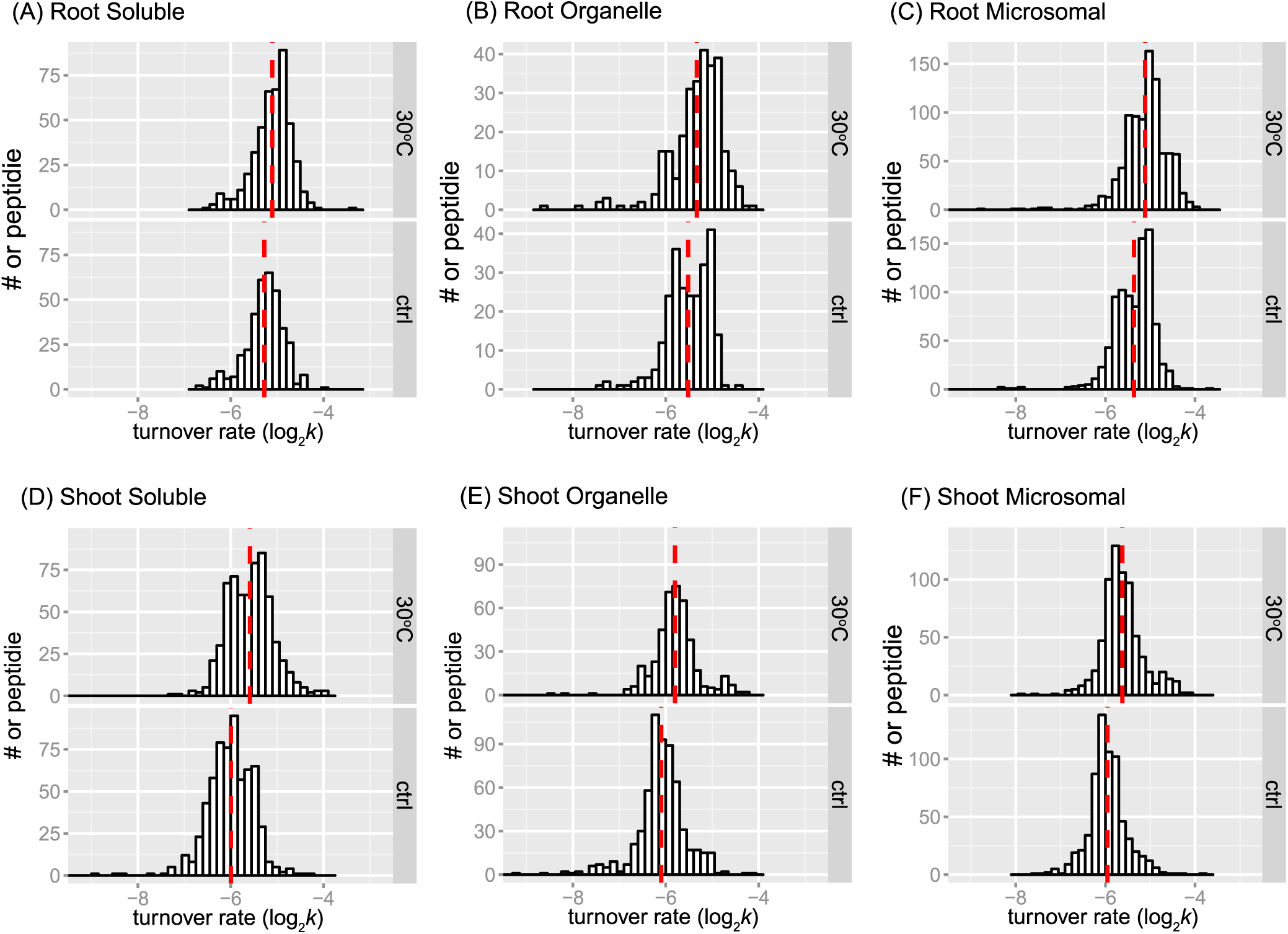
Peptide turnover rate distributions by tissue, fraction, and treatment. Histograms show peptide log_2_*k* values plotted for enriched soluble, organelle, and microsomal fractions of root (panel A, B, C) or shoot (panel D, E, and F) tissues. The control (ctrl) and 30°C groups are plotted in the bottom and top frame, respectively. The *y*-axis is the number of peptide counts. The mean value is plotted as dashed line in red. The bin width is 0.15 for all histograms.

**Figure 3.**
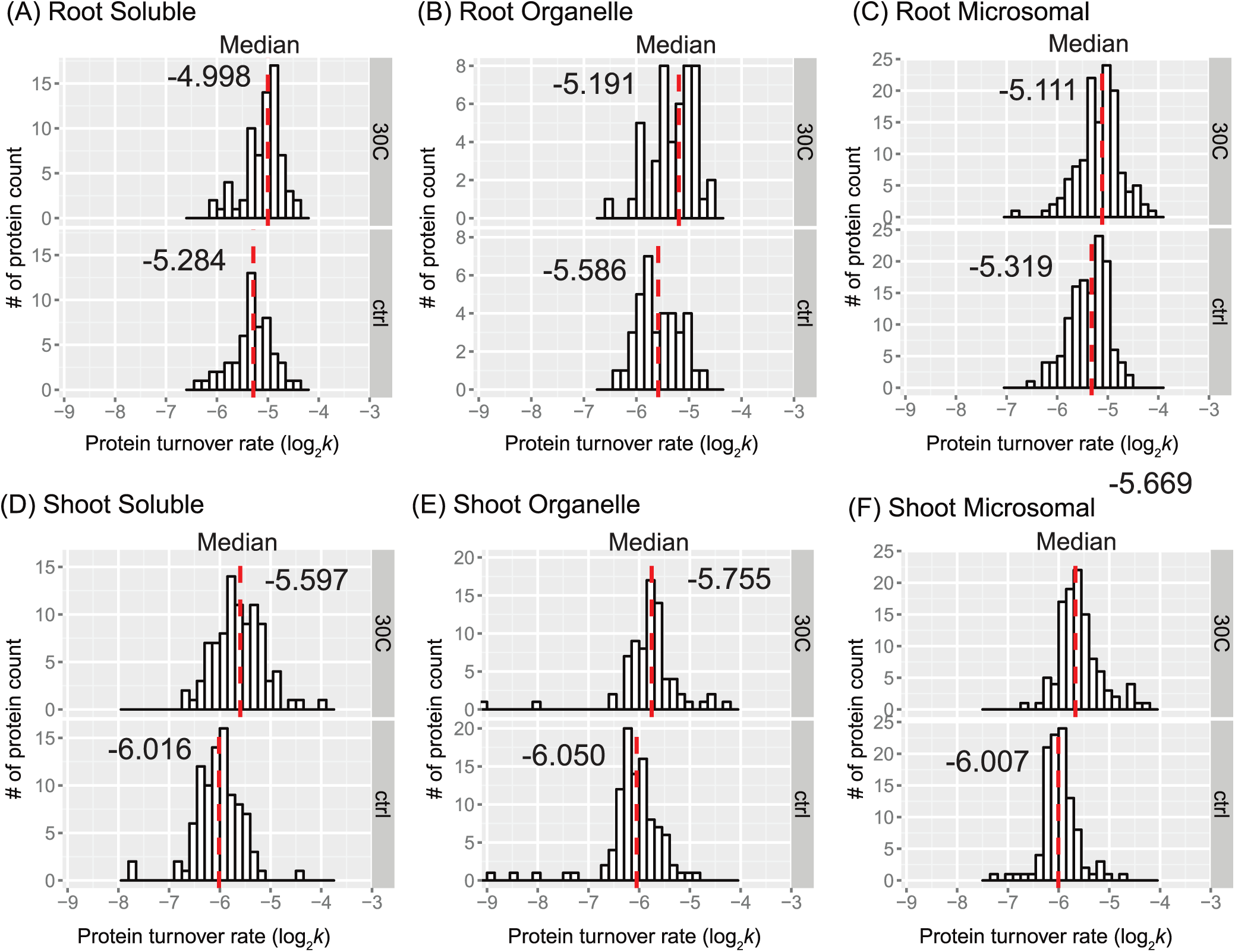
Protein turnover rate distributions by tissue, fraction, and treatment. Histograms show protein log_2_*k* values plotted for enriched soluble, organelle, and microsomal fraction of root (panel A, B, and C) or shoot (panel D, E, and F) tissues. The control (ctrl) and 30°C group is plotted in the bottom and top frame, respectively, The *y*-axis is the number of protein counts. The median value is labeled and plotted as dashed line in red. The bin width is 0.15 for all histograms.

As the mean may be a more robust population estimator than the median for the bimodal distribution, the mean value was shown in each peptide rate distribution in Figure 2. In every fraction of root or shoot tissue, the average of peptide log_2_*k* of the 30°C group was less than that of the control, indicating that peptides tend to turn over faster in response to higher temperature. The difference in the mean of log_2_*k* between the control and 30°C was about 0.17 in the root enriched soluble fraction, 0.18 in the root organelle enriched fraction, 0.25 in the root microsomal enriched fraction, 0.41 in the shoot soluble enriched fraction, 0.30 in the shoot organelle enriched fraction, and 0.33 in the shoot microsomal enriched fraction. Therefore, there was a 1.12∼1.18-fold change in turnover rate of root peptides and a 1.23∼1.32-fold change in turnover rate of shoot peptides at elevated temperature. At the level of proteins, the fold change of average turnover rate due to 30°C stress ranged from 1.16 in the root enriched soluble fraction, ∼1.31 in the root organelle enriched fraction, 1.22 in the root microsomal enriched fraction, 1.26 in the shoot soluble enriched fraction, 1.23 in the shoot organelle enriched fraction, and 1.34 in the shoot microsomal enriched fraction. Both peptide and protein turnover rate distributions in the three protein fractions indicate that shoot and root proteomes have different scales of response to high temperature. Comparing the change in protein turnover rate between roots and shoots in response to high temperature using ANOVA and Tukey’s HSD test revealed a significant difference in log_2_*k* (p < 0.001).

The histograms of some data groups exhibit bell-shaped distributions with slightly asymmetrical patterns in both control and treatment groups. It is possible that the bimodality at the peptide level reflects variations in amino acid content, which could influence peptide turnover rate calculations. In general, the presence of bimodality is less apparent in the protein turnover histograms (Figure 3) compared to the peptide histograms (Figure 2). This observation is not surprising given the significant decrease in the number of observations from peptides to protein turnover. One potential method to test for bimodality is by employing Hartigan’s dip test [29]. In the dip test, the null hypothesis states that the distribution of the sample is unimodal, while the alternative hypothesis suggests that the distribution is not unimodal, indicating at least bimodality. The results from the dip test indicated significant non-unimodal or at least bimodal distribution of peptide turnover rate (*k*) in the control group of the root microsomal fraction (p-value = 0.00376) and marginally non-unimodal in the root organellar fraction (p-value = 0.0847).

#### 2.2.2. Coefficient of variation in protein turnover as a function of the number of peptide observations

Figure 4 shows the extent of variation in protein turnover in this experiment as a function of the number of peptide observations that were averaged to produce the rate for each protein. Since the protein turnover rates were obtained as the mean of turnover rates of all selected peptides, the coefficient of variation (CV), also known as relative standard deviation, can be used to show variability in relation to the mean of the population. Here, the values of CV were calculated as the standard deviation divided by the absolute value of protein turnover rate log_2_*k*. Comparing Figure 4A and 4B, it appears that both the control and 30°C datasets have similar levels of variability, suggesting consistency in the protein turnover rates between these two groups. At first, it appears as though the CV values for the protein turnover rates are larger for the rate values calculated from smaller numbers of peptides, but the median CV ranged from 0.02 to 0.05 and is independent of peptide number. The illusion of high CV for small numbers of peptides is due to the inverse correlation between the numbers of rates calculated and the number of peptides used for each calculation. As a result, there are significantly more real outliers for the very welldefined distribution of CV of protein turnover rates from 2 peptides. Most CV values are within the range of 0 to 0.10, while less than ∼10 proteins have a CV greater than 0.10. When only 2 peptides were computable for one protein, there were only 3 or 4 cases where the CV was greater than 0.15. Given this analysis of CV, it is quite reasonable to include proteins with turnover rates calculated from as few as 2 computable peptides and to make protein turnover rate comparisons between samples with different numbers of computable peptides.

**Figure 4.**
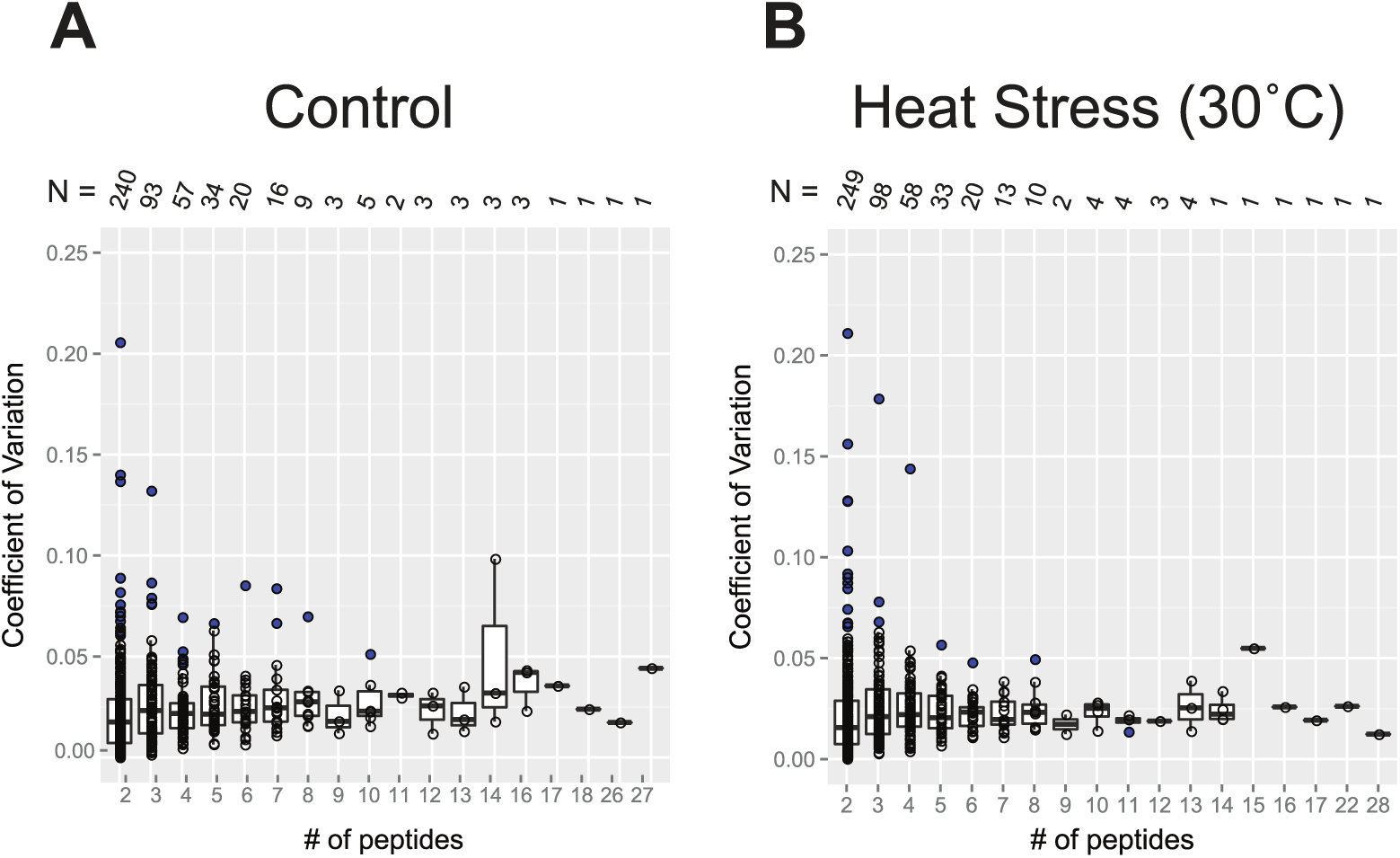
Box plots of the coefficient of variation (CV) of protein turnover rates plotted as a function of the number of peptide rates used in each calculation. The value of CV was calculated from standard deviation of log2k divided by the mean of log2k. The entire dataset used in this plot analysis includes all unique and shared peptidesand is separated based on the treatment group: the control (panel A) and 30°C (panel B). Boxes show the interquartile range (IQR) of turnover rates of proteins. The error bar represents the entire range of rates and the blue dots represent outliers (1.5 IQR). The number of data points in each x-axis category is given as *N*, below the x-axis of both plots.

#### 2.2.3. Statistical significance of changes in protein turnover rates upon heat treatment

Proteomic analysis of protein turnover requires a large number of individual UHPLC-HRMS/MS analyses to provide data across multiple time points, different tissues, different biochemical fractions, and test conditions. These analyses take a considerable amount of time and are expensive. For this reason, it is often impractical to use sampling of biological replicates as a means of testing statistical significance. Furthermore, these analyses often fail to identify many of the lower abundance proteins in replicate runs due to the element of chance in precursor ion detection. As a result, replicated peptide observations are only available for a portion of the identified proteins and typically only those in the top several orders of magnitude in protein abundance. Given the time, cost, and repeatable coverage considerations, a reasonable alternative for determining the significance of changes in turnover rate (log_2_*k*) between treatments is to apply a linear mixed-effect model (LMM) [30]. An LMM allows one to estimate the likelihood of a difference in log_2_*k* values between treatments using a linear model consisting of a mixture of fixed and random effects. The fixed effects represent the errors associated with the conventional linear and non-linear regression portions of the turnover rate derivation, and the random effects represent unknown but random effects such as how peptides were selected from the population of peptides during the UHPLC-HRMS/MS analysis. The LMM approach is also compatible with taking the average of the peptide turnover rate values to determine the protein turnover rate. Supplementary Table S-1 lists the output of the LMM estimation.

Summary of the number of identified peptides and proteins in this study, with applied threshold for selection, and their number with significant changes in turnover rate (log_2_*k*) due to the 30°C treatment (p < 0.05) identified in the enriched soluble, organellar, and microsomal fractions of *Arabidopsis* seedling root or shoot tissues are listed in Table S-1. The identified proteins with significant changes in turnover rate (log_2_*k*) are listed in Supplementary Table S-1, with at least 1 unique peptide in both control and 30°C samples, which were discussed further (Figures 5, 7, and 8). An overview of the distributions of estimated differences in protein turnover rates between control and heat stress is shown in Figure 5 as histograms (Figure 5A) or box plots (Figure 5B). Overall, proteins enriched in the shoot soluble fraction had the largest change in turnover rate with a median increase of ∼0.492 log base 2 scale, or ∼1.41-fold increase in protein turnover rate (*k*) upon heat stress. The box plots in Figure 5B demonstrate that all but the root or shoot soluble fraction had similar variation in the change of protein turnover rate upon heat stress. ANOVA and Tukey’s HSD tests revealed that there was a significant difference in the fold change of turnover rate between root and shoot soluble fractions (p < 0.001). There were also differences between shoot soluble and shoot organellar fractions (p < 0.01) and root soluble and root microsomal fractions (p < 0.05). It suggests that the proteins in the shoot tissue exhibit a greater change in rates of turnover in response to high temperature than proteins in the root tissue. Hence, the root proteome may not be as responsive as the shoot proteome to temperature change.

**Figure 5.**
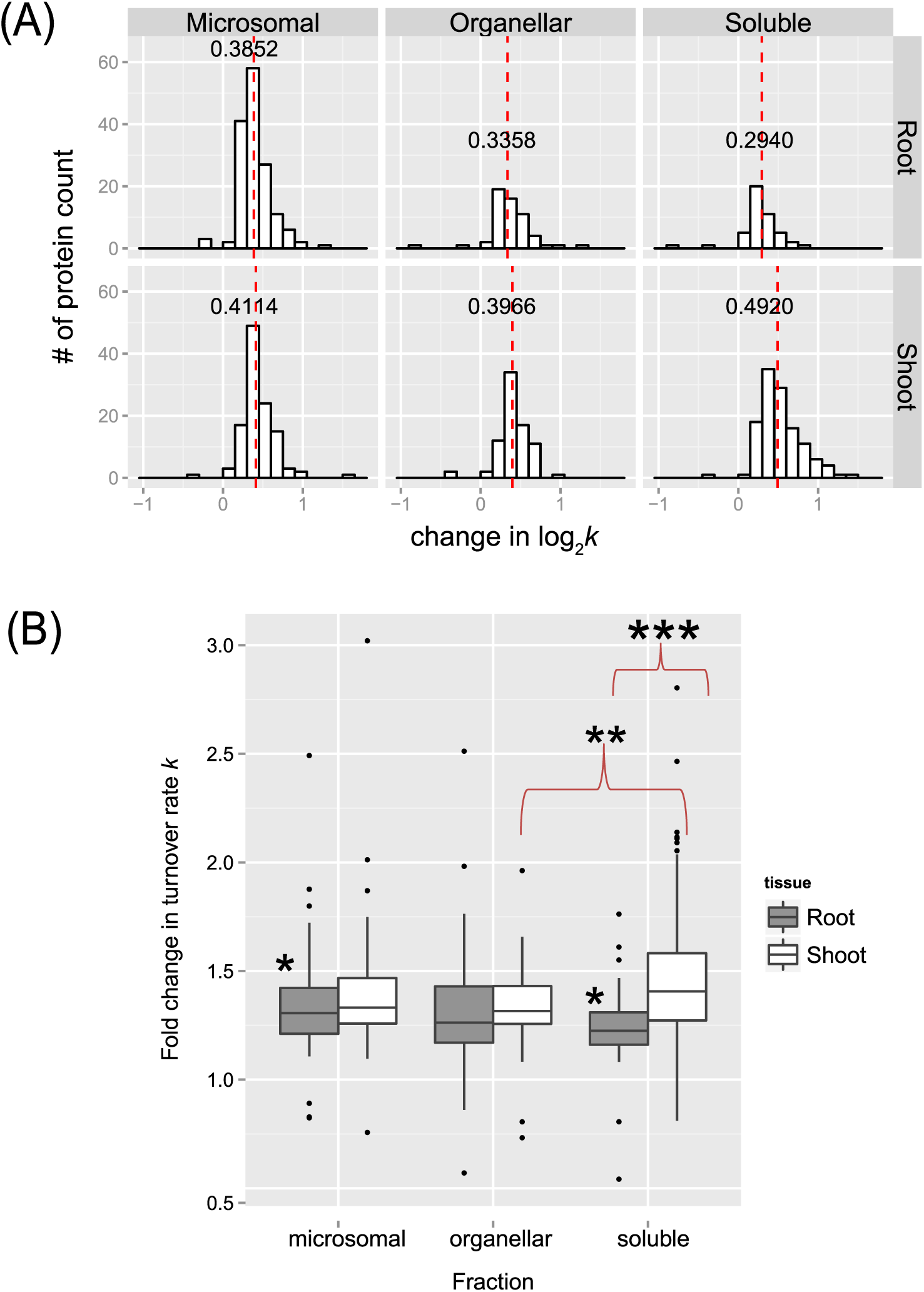
The distribution of changes in protein turnover rates across different tissue and enriched fractions. (A) Histograms showing distributions in the estimated fold change in protein turnover rate*k* values in response to 30°C plotted for soluble, organelle, and microsomal fraction of roots (on the top) or shoots (on the bottom), respectively. The bin width is 0.15 for all histograms. The median value is labeled and plotted as dashed line in red. (B) Box plots of estimated estimated fold change in protein turnover rate(*k*) in response to 30°C of protein identified in the root and shoot enriched soluble, organelle, and microsomal fractions. The analyzed data only include proteins with significant change in log_2_*k* (p <0.05) at least 1 unique peptides in both control and 30°C group, which was estimated using a LMM approach after peptide selection criteria were applied. Boxes show the interquartile range (IQR) of change in turnover rates *k*. The error bar represents the entire range of rates and the closed circles represent outliers (1.5 IQR). The estimated change in turnover rates were analyzed by Tukey’s HSD (honest significant difference) test and * for p < 0.05, ** for p < 0.01, *** for p < 0.001.

### 2.3. Links Between Protein Functional Categories and Changes in Protein Turnover Rates Upon Heat Treatment

#### 2.3.1. Protein function and turnover rates of proteins

In a comparison of shoot and root soluble fractions, the proteins in shoots exhibited a much higher change in turnover rates than in the roots (Figure 5). To determine if function might play a role in protein stability, root and shoot proteins in enriched soluble and membrane fractions from the control experiment were sorted into functional categories. The functional categories were adapted from the MapCave website using the TAIR10 database. Shown in Figure 6 are box and whisker plots of turnover rates of root (panel A) and shoot (panel B) proteins from the control experiment categorized by functional groups. Only proteins with at least 2 unique peptides were reported in Figure 6. For groups with at least 3 proteins, most of them had fairly similar variation in log_2_*k* values, such as glutathione S-transferase (GST), protein synthesis, protein targeting, glycolysis, mitochondrial electron chain/ATP synthesis, cellular transport, and stress in root proteins or amino acid metabolism, the light reaction of photosynthesis, the Calvin cycle of photosynthesis, and protein folding in shoot proteins as these categories have well-studied proteins with known function. Some proteins appeared to have more variation in log_2_*k* values, especially the ones in the functional categories like redox reaction (ranged from −4.97 to −6.17 in roots, −4.48 to −6.81 in shoots), signaling (−4.89 to −6.12 in roots), development (−4.96 to −6.24 in roots), or secondary metabolism (−4.74 to −7.44 in shoots) as the proteins in these groups are involved in more varieties of function.

**Figure 6.**
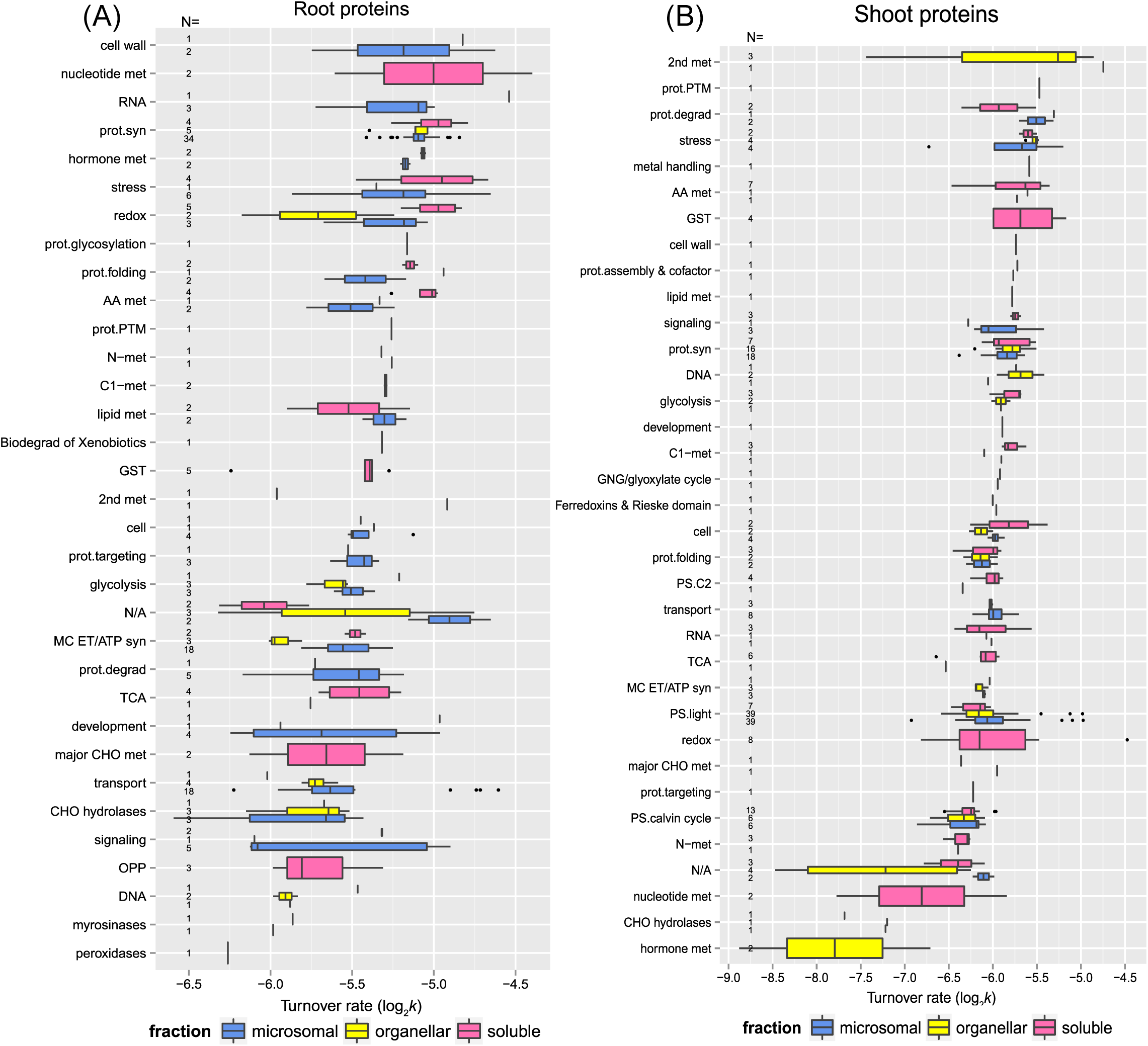
The relationship between protein function and protein turnover rates. Box plots of protein turnover rate log_2_*k* for root (Panel A) and shoot (Panel B) proteins from the control experiment are sorted by functional categorization, which was adapted from the MapCave website (http://mapman.gabipd.org/web/guest/mapcave) using TAIR10 database, with outliers shown as closed circles. The used data only include proteins at least 2 unique peptides. The number of protein in each function category is given as *N*, along the y-axis of both plots. The protein count of each function group is also labeled in the plot. Abbreviations: 2nd met, secondary metabolism; AA met, amino acid metabolism, C1-met, single carbon metabolism; DNA, CHO hydrolases, miscellaneous gluco-, galacto- and mannosidases; DNA processing; Glc-, Gal& mannosidases, glucosyl-, galactosyl& mannosylglycohydrolases; GNG, gluconeogenesis; GST, glutathione S-transferases; lipid met, lipid metabolism; major CHO met, major carbohydrate metabolism; MIP, major-intrinsic proteins; MC. ET/ATP syn, mitochondrial electron transport/ATP synthesis; N-met, nitrogen metabolism; OPP, oxidative pentose phosphate pathway; prot.assembly, protein assembly & cofactor ligation; prot.degrad, protein degradation; prot.folding; protein folding; prot.targeting, protein targeting; prot.PTM, protein post-translational modification; prot.syn, protein synthesis; PS.C2, photorespiration; PS.light, the light reaction of photosynthesis; PS.calvin cycle; the Calvin Cyle of photosynthesis; RNA, RNA processing; S-assimilation, sulfur assimilation; TCA, tricarboxylic acid cycle.

Some functional categories exhibited somewhat faster turnover rates, as shown by higher median log_2_*k* values in Figure 6. It has been believed that proteins with faster turnover rates could be potential control and regulation points as Heat Shock Proteins (HSPs), proteins involved in signaling, protein synthesis and degradation, and DNA/RNA processing enzymes turned over faster in the proteome turnover study using *Arabidopsis* cell culture; while glycolytic enzymes had the slowest turnover rates. The protein function seem to be related to turnover rates in general in this study. For example, the root proteins involved in cell wall formation, nucleotide metabolism, RNA processes, protein synthesis, hormone metabolism, and stress response had faster turnover rates; while proteins involved in DNA processes, oxidative pentose phosphate pathway, major carbohydrate metabolism, and signaling had slower turnover rates. In the shoot tissue, proteins related to secondary metabolism, protein degradation, stress response had higher turnover rates appeared to turnover faster, while proteins involved in the Calvin cycle, hormone, and nucleotide metabolism had much lower turnover rates.

Some specific proteins and their turnover rates were of special interest. Table 1.1 and 2.2 listed the top 10 fastest and slowest proteins in the control experiment of root and shoot tissues, respectively. As listed in Supplemental Table S-2, there was a 4.58-fold difference between the lowest to the highest turnover rate (*k*) among the identified root proteins (total number 221) while there was a 21.12-fold difference between the lowest to the highest in turnover rate among the shoot proteins (total number 297). Therefore, the root proteome appeared to turnover faster but with less variation in general, which suggests there might be a closer correlation in regulating protein synthesis and degradation in root tissue. Stress or redox signaling-related proteins like HSP 70-1 and Chaperone protein dnaJ 3 in roots or HSP 70-11 and Catalase-3 in shoots exhibited relatively rapid turnover. Proteins involved in the light reaction of photosynthesis, especially Photosystem II D2 protein and Photosystem II CP43 reaction center protein, turned over much faster than other proteins functioning in photosynthesis. Therefore, these two proteins might need to be replaced rapidly to maintain normal carbon fixation in plants. Some transport proteins like plasma membrane ATPase 1 (AHA1) and ABC transporter G family member 36 (ABCG36; PEN3; PDR8) in the root tissue were identified as outliers in the box plot due to their extraordinarily fast turnover rates. It has been shown that the expression of ABCG36/PEN3/PDR8 gene in seedlings is 5 to 40 fold higher than other ABC transporters and its transcript abundance in leaves is comparable with transcript levels of some house-keeping genes like cytosolic glyceraldehyde-3-phosphate, suggesting the multiple physiological functions of ABCG36/PEN3/PDR8. It has later been reported that ABCG36/PEN3/PDR8 is an ATP-binding cassette (ABC) transporter localized on the plasma membrane and is thought to efflux indole-3-butyric acid (IBA) in root tips, several biotic, and abiotic stress responses. The fast turnover rate of ABCG36/PEN3/PDR8 in seedling roots could result from the high level of protein synthesis, supporting its multiple roles in heavy metal ion tolerance as well as regulating the IBA-mediated homeostasis of auxin in roots. On the other hand, some glycosyl hydrolase family proteins, such as betaglucosidase 22 (BGLU22) or beta-glucosidase 23 (BGLU23/PYK10) in the root or shoot tissue had the slowest turnover rates. BGLU family proteins are important for ER formation and their hydrolytic activity for glucoside that accumulates in the roots of *Arabidopsis* has been believed to be important in defense against pests and fungi. It has been proposed that healthy seedling roots accumulate beta-glucosidases in the ER bodies. Therefore, when plant cells are under attack from herbivore or pathogen, beta-glucosidases would leak from the ER body and bind to GDSL lipase-like proteins (GLLs) and Jacalin-related lectins in the cytosol to form complexes with increased enzyme activity which hydrolyzes glucosides to produce toxic compounds like scopolin. These proteins are very abundant and expressed exclusively in *Arabidopsis* seedlings, so their slowest turnover rates identified in this study suggest that BGLU22 and BGLU23 act like housekeeping proteins in *Arabidopsis* seedlings in order to rapidly trigger defense mechanism on demand.

**Table 1.1.**
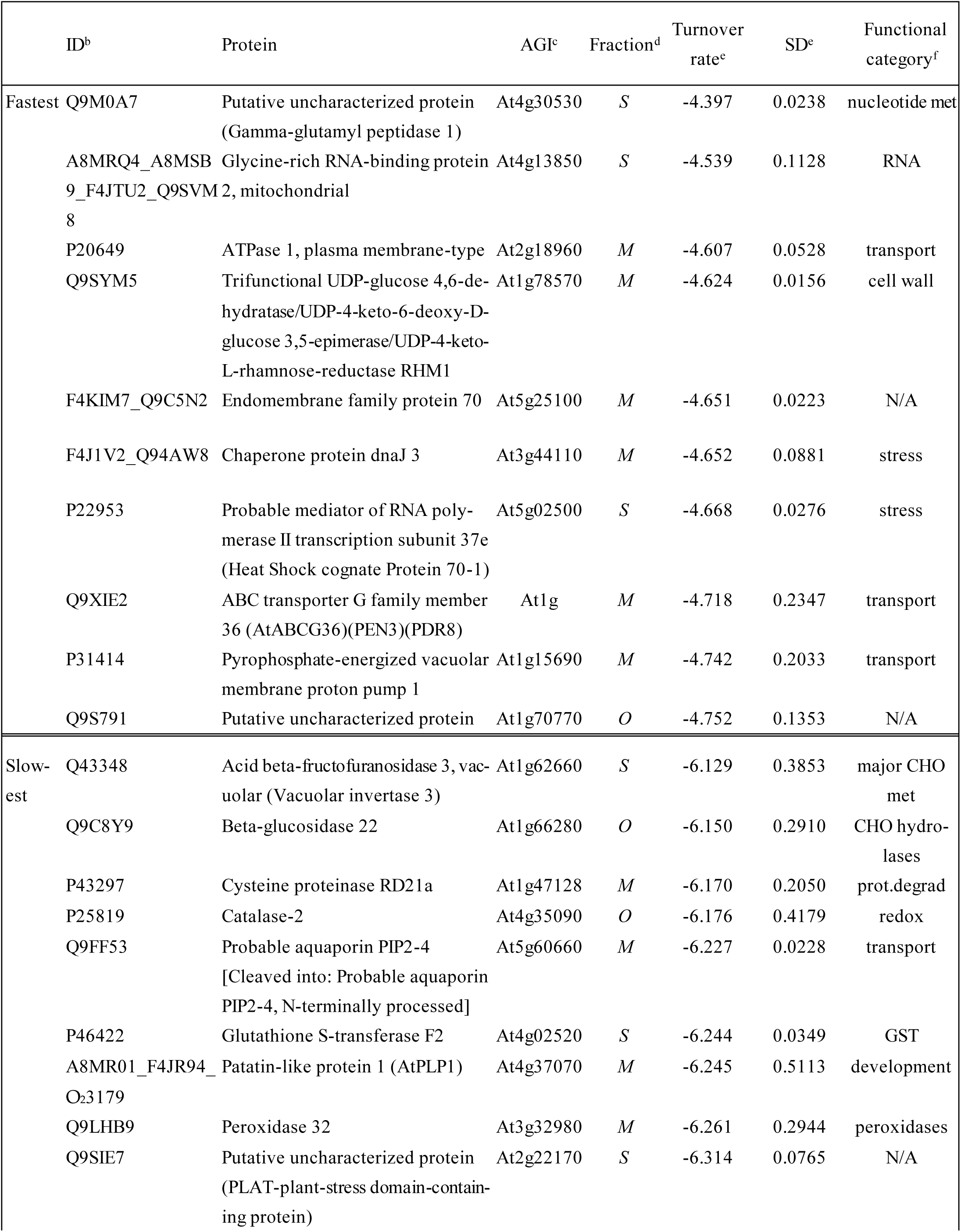
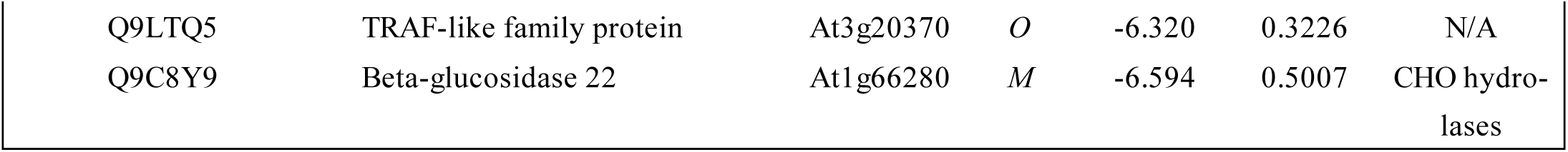
The 10 fastest and lowest turnover proteins in the enriched soluble or membrane fraction of *Arabidopsis* root**s^a^**.

**Table 1.2.**
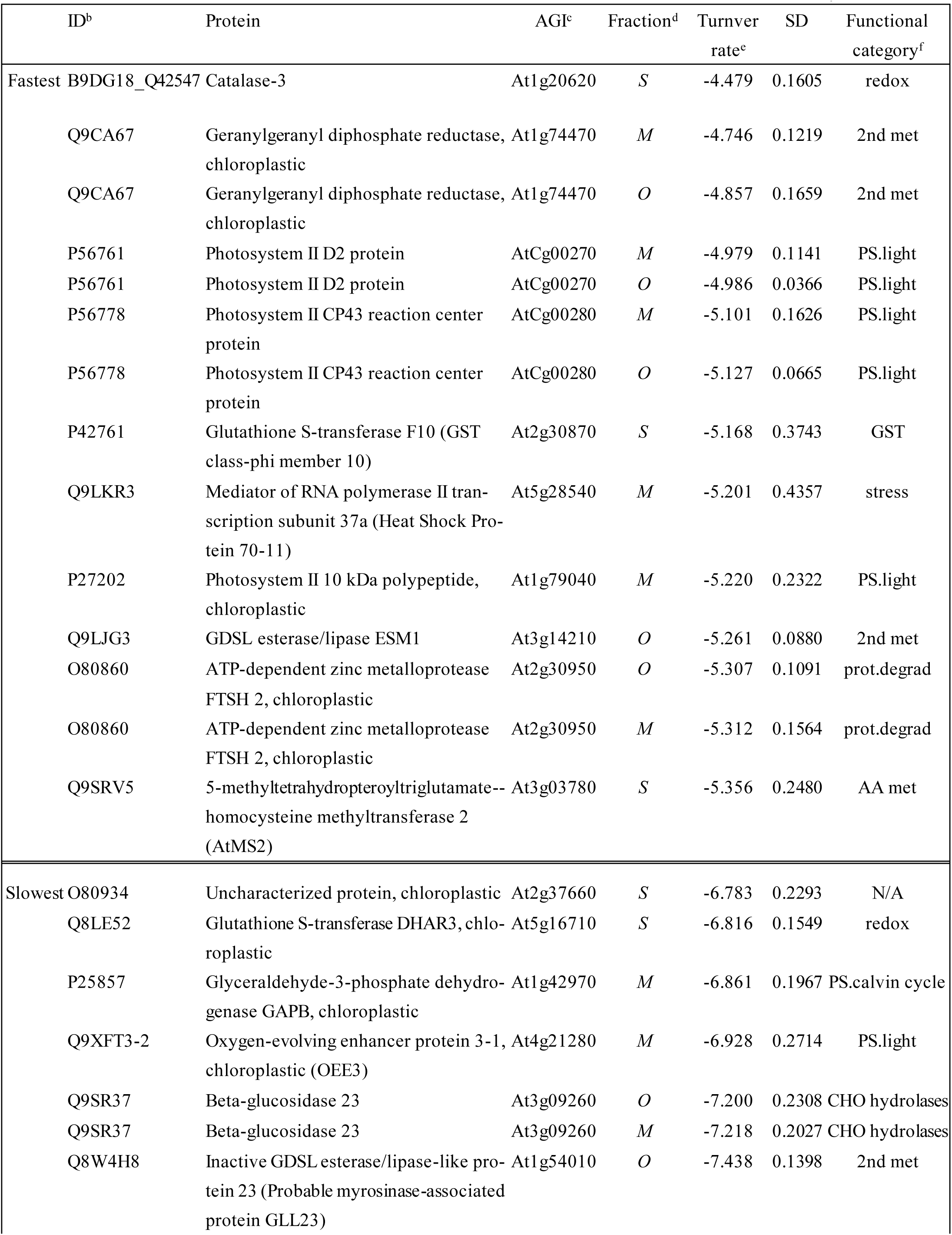
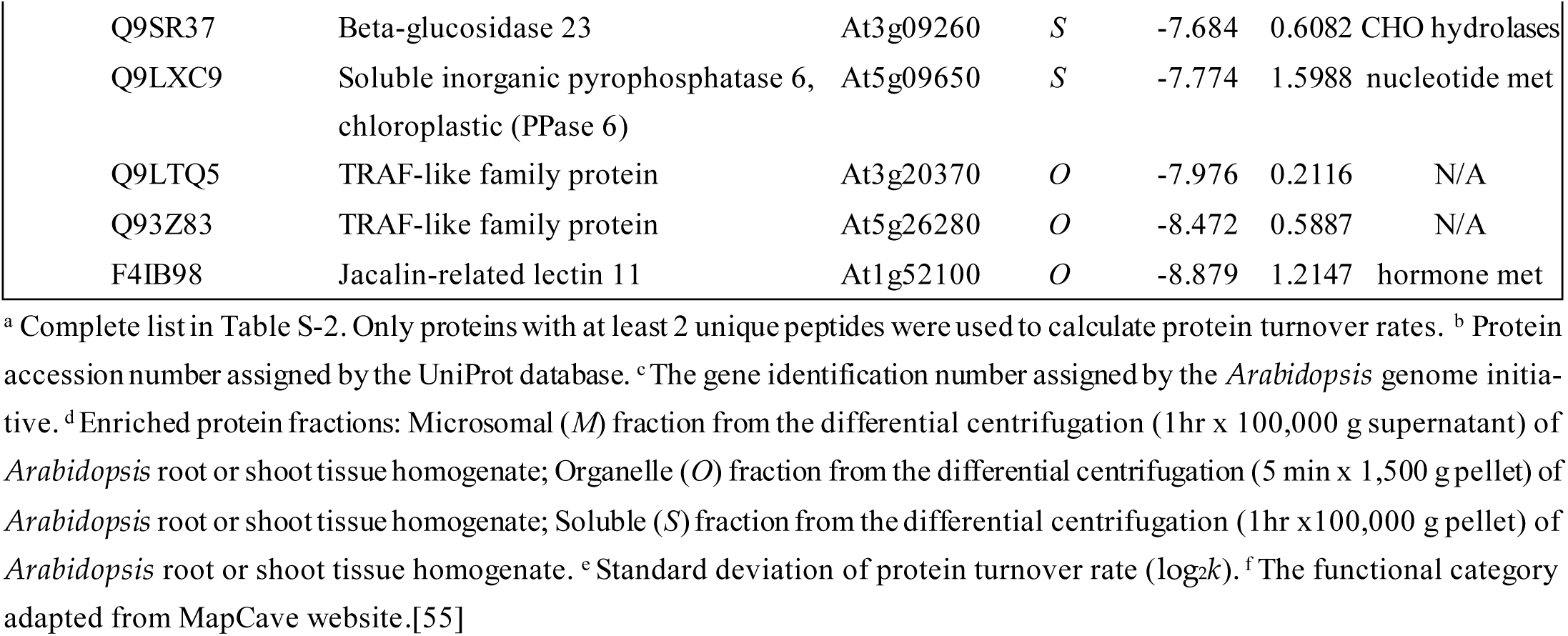
The 10 fastest and slowest turnover proteins in enriched soluble or membrane fraction of *Arabidopsis* shoots ^a^.

#### 2.3.2. Protein function and change in turnover rates due to high temperature

To further explore functional correlations with protein turnover changes during heat stress, the proteins with significant changes due to high temperature identified in this study were also sorted into functional categories. Figure 7A and B are box plots showing the fold changes in turnover rate in response to high temperature treatment (calculated from the estimated difference in log_2_*k* between the control and 30°C using the LMM fit) across functional categories for each tissue and fraction. Only proteins with a significant change in log_2_*k* (p < 0.05) and at least 1 unique peptide in both control and 30°C groups were included in this analysis. In each plot, the protein categories were sorted on the y-axis from largest to smallest median difference in protein log_2_*k*. Functional categories with only 1 data point (1 protein) were included in the plot to provide additional coverage of the functional categories. The number of proteins in each functional category is given as N along the y-axis of each plot. Most of the groups had median values ranging from 1.25 to 1.75 fold change. Among those identified in roots, proteins involved in redox signaling pathways, stress response, protein folding, and calcium-signaling pathways had the largest median changes in turnover rate. In shoots, the beta-glucosidase family and proteins sorted in photorespiration, protein folding, stress response, hormone, and secondary metabolism exhibited the largest median changes in turnover rate due to heat (∼1.5 fold change in *k*).

**Figure 7.**
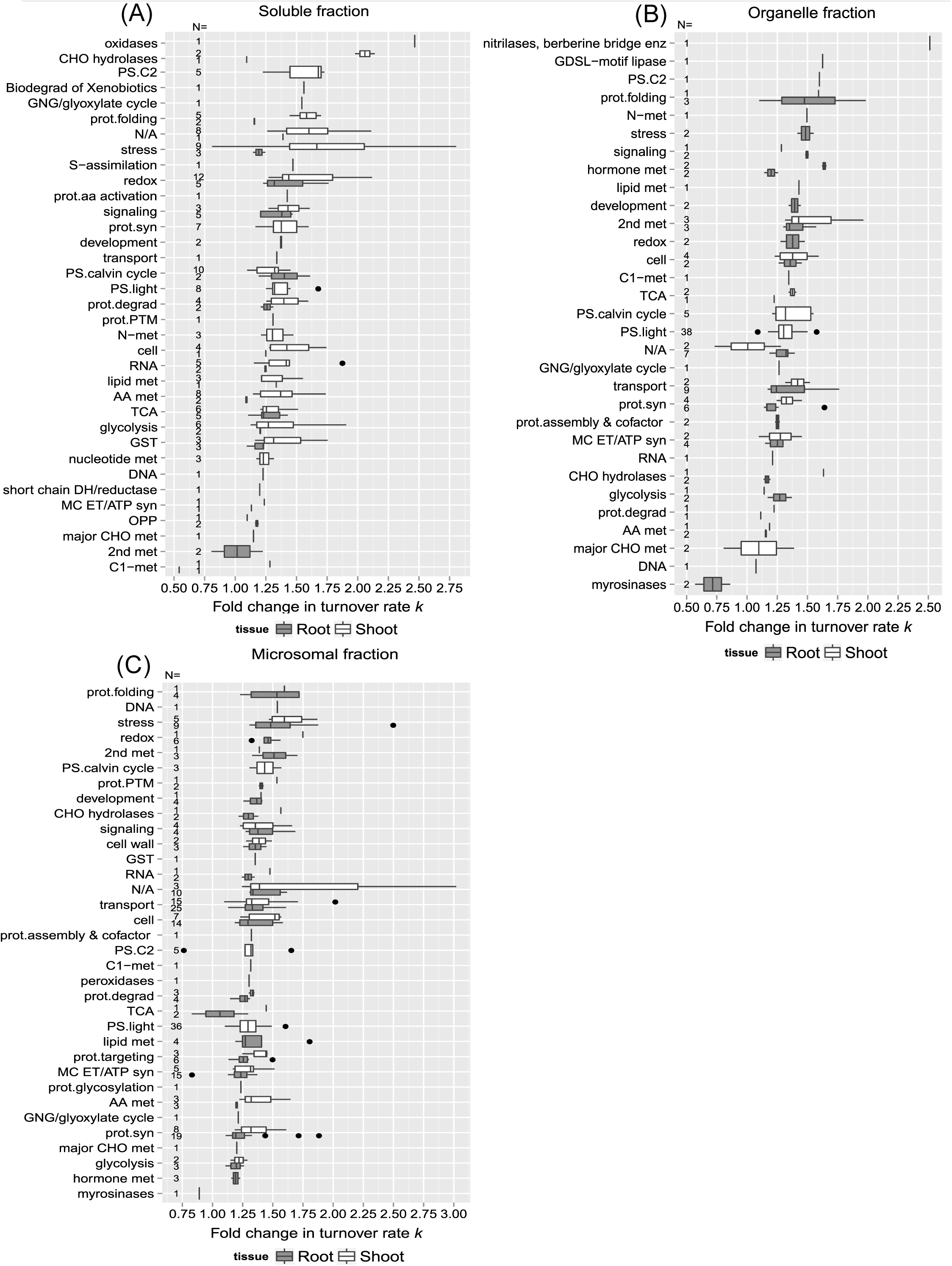
The relationship between protein function and change in turnover due to elevated temperature. Box plots of estimated change in log2k in response to 30°C (diffs. in log2k) for root and shoot proteins are sorted by functional categorization, which was adapted from the MapCave website using TAIR10 database, with outliers shown as closed circles. (A) Soluble enriched fraction of root or shoot tissue homogenate. (B) Organelle enriched fraction of root or shoot tissue homogenate. (C) Microsomal enriched protein fraction of root or shoot tissue homogenate. The used data only include proteins with significant change in log2k (p <0.05) and at least 1 unique peptides in both control and 30°C group. The number of protein in each function category is given as N, along the y-axis of all plots.

In the functional categories identified in both root and shoot soluble fractions such as redox signaling, stress response, protein degradation, and glutathione S-transferase metabolism, shoot proteins exhibited greater changes in turnover rates than root proteins, as well as secondary metabolism, protein synthesis, and stress response in the enriched organellar and microsomal fractions (Figures 7B & 7C). On the other hand, proteins assigned to the glycolysis, cellular transport, mitochondrial electron chain/ATP synthesis functional group, TCA cycle, signaling, cell organization, and cell wall structure displayed similar changes in turnover rate with heat stress in both roots and shoots, suggesting that the turnover of proteins involved in these biological processes such as mitochondrial ATP synthesis is regulated uniformly throughout the whole seedling.

Comparing the changes in turnover rates of proteins within the same functional category between different root (Figure 8A) or shoot (Figure 8B) fractions could help identify specific proteins with different levels of responses to heat stress due to compartmentalization. For example, shoot proteins involved in photorespiration appeared to be more affected by high temperature in the soluble fraction than in the membrane fractions in general. Although after inspection of the proteins listed in Supplementary Table S-1, exceptions such as Glycolate oxidase 1 (GOX 1; At3g14420), which was identified in both soluble and microsomal fractions, turned over fairly rapidly in both fractions. Other categories such as the light reaction of photosynthesis, cellular transport, cellular organization, mitochondrial electron transfer/ATP synthesis, protein synthesis, and glycolysis exhibited a similar breadth of responses across different fractions. This may be due to the fact that these proteins were relatively abundant so that they are being isolated in multiple fractions. Choroplastic ATP synthase subunit alpha (Atcg00120), for example, was identified in all three fractions.

**Figure 8.**
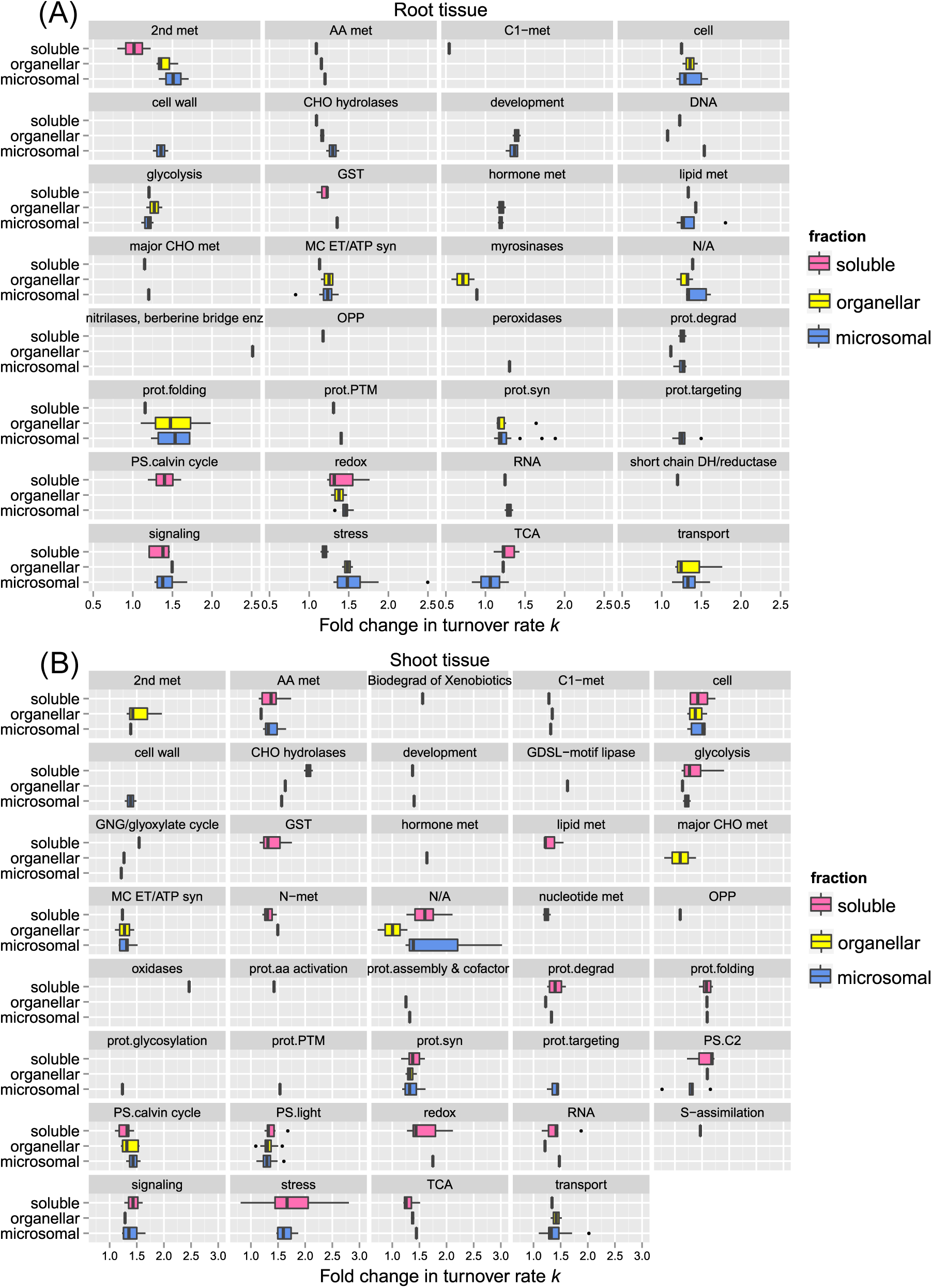
Comparison of protein functions with the change of turnover rates in response to 30°C between different protein fractions. Boxes show the interquartile range (IQR) of estimated difference in log_2_*k* turnover rates of proteins (diffs in log_2_*k*). Proteins are sorted in functional categorization, comparing results between the enriched soluble, organelle, and microsomal fraction of root (panel A) or shoot (panel B) tissues. The error bar represents the entire range of rates and the closed circles represent outliers (1.5 IQR).

## 3. Discussion

In this study, a high temperature treatment of 30°C was applied for durations ranging from 8 to 48 hours. Despite being relatively moderate compared to typical heat stress studies, it has been demonstrated that even a modest change in temperature, such as transferring 12-day-old *Arabidopsis* seedlings from 12 to 27°C for 2 hours, can significantly alter the expression of over 5000 genes by at least 2-fold [31]. The present study’s moderate high temperature treatment aligns with the moderately elevated temperature, contrasting with the heat stress conditions in the Mittler study [31]. This suggests that different heat sensors and signaling pathways may perceive these temperature regimes differently.

Results of this study suggest that heat stress cause greater change in shoot proteome than root proteome. In plants, it is believed that root growth is more sensitive to acute heat stress than shoot growth as high soil temperature is more detrimental than high air temperature, and lower soil temperature could help plants survive when grown at high air temperature [32]. Future studies may employ metabolic flux analysis [33–35] to delve into metabolite turnover in response to high temperature treatments, offering insights into molecular-level plant adaptation and aiding the development of strategies to enhance crop heat tolerance and mitigate climate change’s agricultural impact.

### 3.1. Heat shock proteins (HSPs) and chaperones

It has long been known that the expression of stress proteins like HSPs could be induced by heat shock at almost all stages of development, and the induction of HSPs seems to be a universal response to heat stress among organisms [36]. In the results, HSPs appear in the stress protein functional category(Table 5). While it is clear that most of the proteins listed in the table are specifically related to heat stress, such as HSP70 −1, HSP70-3, HSP70-11, HSP90-2, HSP90-3, and the chaperone protein htpG family, in several of the fractions, there are additional potential stress response-related proteins predicted from the microarray gene expression data, such as RD2 protein (involved in the response to desiccation), major latex protein (MLP)-like proteins 328 and 34 (responsive to biotic stimulus), MLP-like protein 34, Dehydrin COR47 (responsive to cold), and At4g23670 protein (involved in the response to salt stress and bacterial infections) [37]. Interestingly, the root soluble fraction HSPs and stress-related proteins had smaller increases in turnover rate compared with other fractions. This significantly smaller increase in HSP and stress-related protein turnover for the root soluble protein fraction may help explain the generally much smaller change in turnover rate in that fraction compared with the other fractions.

A previous study found that stress response proteins such as heat shock chaperones and proteins associated with oxidative stress have relatively high degradation rates, although that study was performed using an enriched mitochondrial fraction of *Arabidopsis* suspension cells [26]. While it is risky to extrapolate from this prior study to intact plants, it is reasonable to postulate that the rapid turnover rate could be even more dramatic in planta. As HSPs help to prevent protein degradation and that the aggregation of HSPs into a granular structure in the cytoplasm helps to protect the protein biosynthesis machineryfrom denaturation [5]. Our study indicates the shoot HSP90-5 (Chaperone Protein htpG family Protein; At2g04030) had a 2.05 fold increase in *k* in response to heat. Overexpression of HSP90-5 in *Arabidopsis* has been shown to result in reduced plant tolerance to drought, salt, and oxidative stress, while knocking out the HSP90 −5 gene results in an embryo lethal phenotype, indicating that HSP90-5 is an essential gene [38]. It has been shown that HSP90-5 is important in maintaining the integrity of chloroplast thylakoid formation [38]. These findings, along with the dramatic change in turnover rate of HSP90 −5 when treated with high temperature in this study all suggest that properly controlled expression of HSP90-5 is important for plant growth and chloroplast biogenesis. HSPs like HSP70, HSP90, and HSP60 belong to molecular chaperone families. Molecular chaperones bind and catalytically unfold misfolded and aggregated proteins as a primary cellular defensive and housekeeping function [39]. Other proteins with significant changes in turnover rate in response to high temperature are also involved in protein degradation and protein folding functions, including several proteinases and multiple chaperones ( Supplementary Table S-1 & Figure 7), including mitochondrial and chloroplast Chaperonin *CPN60* (HSP60) and CPN-10, which turnover rapidly in response to heat. Plastidic *CPN60* alpha and beta are required for plastid division in *Arabidopsis* and *CPN60* are required to be maintained at a proper level for folding of stromal plastid division proteins and are essential for development in chloroplasts [40]. The observed change in *CPN60* turnover rates is somewhat correlated to the study revealing the slightly reduced expression of CPN-60 in seedling shoots when encountering the elevated temperature at 28°C [40]. Another chaperone protein AtBAG7 (At5g62390) exhibited faster turnover rate at elevated temperature. AtBAG7 is required to maintain the c and is localized in the endoplasmic reticulum, which is unique among BAG family members[13,41]. It has been proposed that cactivity may be regulated post-translationally, given that its gene expression does not appear to be affected by heat or cold stresses [13]. Since AtBAG7 directly interacts with an HSP70 paralog, AtBAG7 activity is likely regulated post-translationally through modulation of protein turnover [13].

### 3.2. Photosynthesis and carbon assimilation

As temperature is a crucial factor affecting photosynthetic activity in plants, as expected, proteins involved in photosynthesis, including components of photosystems I & II (PSI & PSII), the cytochrome b6-f complex, chloroplast ATP synthase, and the Calvin cycle, were identified for having varying degrees of change in turnover in response to heat. Prior heat stress-related studies found that the oxygen-evolution complex (OEC) of PSII is the main target of heat stress [42]. From this study, changes in turnover rates of OEC subunits were around 1.21-1.42 fold, similar to the majority of the proteins involved in photosynthesis, in response to heat (Supplementary Table S-1). There were extreme cases like RuBisCO activase (At2g39730) and chlorophyll a/b binding protein (LHCB6; At1g15820) that exhibited larger, 1.57 and 1.60 fold changes in *k*, respectively. As it is highly sensitive to heat denaturation, RubisCo activase is thought to be a key element involved in mediating the heat-dependent regulation of carbon assimilation as it could limit the photosynthetic potential of plant tissues at high temperature [11]. Although the enzyme activity of RubisCo activase was not decreased until the temperature was higher than 37°C in cotton and tomato leaves [11], our study suggests that this enzyme in *Arabidopsis* seedlings could “sense” relatively mild elevated temperatures like 30°C in terms of protein turnover. It is hard to judge from the results whether the turnover rates of proteins of PSII and light-harvesting complex II (LHCII) were more affected by high temperature than PSI, as it has long been believed that PSII is more vulnerable to elevated temperature [43,44]. A comparison, however, of the differences between PSI and PSII protein turnover following heat stress should indicate the relative heat tolerance of the two photosystems under mild elevated temperature conditions. To this end, LHCB6, which is associated with PSII, turns over significantly faster (1.60 fold change in *k*) after heat treatment than the Photosystem I reaction center subunit III (1.25 fold change in *k*). Notably, these rate changes are on the high and low extremes of the range of changes observed for protein components of photosynthesis. LHCB6 is a monomeric antenna protein of PSII, participating in zeaxanthin-dependent photoprotective mechanisms, and is therefore thought to be specialized in enhancing photoprotection under excess light conditions. The presence of the protein is often associated with the adaptation of plants to terrestrial ecosystems [45]. Heat stress at temperatures around 38-40°C has been demonstrated to cause structural changes in the thylakoid membranes, as well as increased phosphorylation of LHCIIs and PSII core subunits, migration of phosphorylated LHCII from the grana stacks to the stroma lamellae, and cyclic electron flow within PSI [46]. It will be interesting to study if the change in LHCB6 turnover could be related to the above observations at 40°C even when more mild temperature conditions like 30°C are employed.

### 3.3. Redox homeostasis: HSPs, catalases and peroxidases

The turnover rates of proteins involved in the production of reactive oxygen species (ROS) were also affected by high temperature. These include several different types of HSPs, catalases and peroxidases. An additional group of antioxidant enzymes, including GST, DHAR, and thioredoxins, exhibited significant heat-related changes in turnover (Supplementary Table S-1). Among those, GST class Tau-member 19 (GSTU19; At1g78380), the most abundant GST in *Arabidopsis*, exhibited the smallest difference in turnover rate (1.31 fold change) in roots but showed a much larger difference (1.75 fold change) in turnover rate in shoots.

Hydrogen peroxide (H_2_O_2_) is an important signaling molecule in plant environmental responses, and heat shock-induced H_2_O_2_ accumulation is required for efficiently inducing the expression of small HSP and ascorbate peroxidase genes (*APX*1 & *APX*2) [47]. Among several types of H_2_O_2_-metabolizing proteins, catalases are highly active enzymes that do not require cellular reductants as they catalyze the dismutation reaction of two molecules of H_2_O_2_ to generate one molecule of O_2_ and two of H_2_O. A 1.40 fold change in turnover rate *k* was observed for catalase-3 (CAT3; At1g20620) in shoots upon temperature elevation. APXs are also known to be important H_2_O_2_-scavenging enzymes, but they use ascorbate as an electron donor. Their function is tightly linked to ROS signaling pathways and the regulation of cellular ROS levels [47]. In this study, there was a moderate increase in APX1 (At1g07890) turnover rates under heat stress conditions in both root and shoot tissues. *APX1* is expressed in roots, leaves, stems, and many other tissues [48], and mutation in *Arabidopsis APX*1 exhibits increased accumulation of cellular H_2_O_2_ and suppressed growth and development [49]. It has been reported that *APX*1 activity could be partially inhibited in roots through modification by S-denitrosylation in an auxin-dependent manner [50]. APX1 could be an interesting research target to explore the links between nitric oxide (NO), H_2_O_2_, auxin hormone signaling, and heat stress.

### 3.4. Special cases: decreases (negative diff.log_2_k) or major increases in log_2_k in re-sponse to heat

#### 3.4.1. GDSL esterase/lipase family

GDSL esterase/lipase 22 (GLL22; At1g54000) showed slightly reduced turnover rates in both root organellar and microsomal fractions (fold change in *k* about 0.86 and 0.89, respectively), indicating that GLL22 becomes more stable and/or with reduced transcription or translation when transferred to 30°C. It has been proposed that under pathogen or herbivore attack, GLL22 may aggregate with beta-glucosidases (BGLU 21, 22, and 23), and other Jacalin-related lectins (JALs) in the cytosol [51]. It is possible that under temperature stress, GLL22 turns over slower due to being recruited into more stable complexes. The change in turnover rates of the BGLU protein family, on the other hand, had a wide variation across root or shoot protein fractions (1.09 ∼ 2.14 fold change due to heat). BGLU proteins appeared to turn over faster in the shoot than root tissue, thus the turnover rates of BGLUs in shoots could be more affected by heat stress than BGLUs in roots. Similar results were observed for JAL proteins like Jacalin-related lectin 30 (PYK10-binding protein 1; At3g16420), Jacalin-related lectin 33 (JAL 33; At3g16450), and Jacalin-related lectin 34 (JAL 34; At3g16460), whose turnover rates also had a greater change in shoots than roots when under heat stress, suggesting these stress-responsive proteins in shoots may be compromised when plants encounter heat stress.

#### 3.4.1. 14-3-3 & v-, p-type ATPase

It is intriguing to observe signaling proteins like 14-3-3 family proteins and proton pump v- and p-type H+-ATPases with significant changes in turnover rate due to elevated temperature because of their known roles in ABA signaling in response to abiotic stress. Increased H_2_O_2_ production under multiple different abiotic stress conditions has been shown to result in elevated levels of ABA, which may in turn be involved in the induction of the temperature stress response in plants [12]. Plant 14-3-3 family proteins function in a wide range of cellular processes. Two 14-3-3 proteins show fairly large changes in protein turnover in response to heat stress: 14-3-3-like Protein GF14 mu (General regulatory factor 9; At2g42590), and 14-3-3-like Protein GF14 epsilon (General regulatory factor 10; At1g22300) with 1.61 and 1.45 fold changes respectively. It has been discovered that 14-3-3 mu participates in light sensing during early development through phytochrome B signalingand affects the time of transition to flowering via interaction with CONSTANS [52]. As T-DNA mutants of the 14-3-3 mu gene exhibit shorter root lengths and a dramatic increase in the numbers of chloroplasts in the roots [53], it is possible that its difference in heat stress response between root and shoot tissues is related to its role in chloroplast development. On the other hand, the 14-3-3 epsilon protein may be involved in brassinosteroid (BR) signaling, like 14-3-3 lambda protein, as the 14-3-3 epsilon protein has been shown to interact with the BZR1 transcription factor in a yeast-two hybrid screen [54]. Therefore, these proteins involved in signal transduction may be affected by heat stress thus influencing the BR hormone regulation.

## 4. Materials and Methods

### Materials

Distilled, deionized water was prepared with a Barnstead B-pure water system (Thermo Scientific, Waltham, MA). Acetonitrile (CHROMASOLV® Plus for HPLC, ≧99.9%), formic acid (ACS reagent ≧96%), and acetone (CHROMASOLV® Plus for HPLC, ≧99.9%) were obtained from Sigma-Aldrich (St. Louis, MO). Triton X-100 was obtained from ICN Biochemicals Inc. (Ohio, USA). 99 atom% K^15^NO_3_ and 98 atom% Ca(^15^NO_3_)_2_ were obtained from Cambridge Isotopes Laboratories, Inc. (Andover, MA). Sequencing grade modified trypsin was purchased from Promega (Madison, WI). Pierce C18 Spin columns were obtained from Thermo Scientific (Pierce Biotechnology, Thermo Scientific, Rockford, IL). Micro-centrifuge tubes used for the proteomics study in this thesis were “Protein LoBind Tube 1.5 mL”, obtained from Eppendorf AG (Hamgurg, Germany). Nylon filter membranes (mesh opening 20 μm, Cat. #146510) were obtained from Spectrum Laboratories Inc. (Rancho Dominguez, CA).

### Plant Growth and Labeling Conditions

*Arabidopsis thaliana* ecotype Columbia Col-0 was used for all experiments. Seeds were sterilized with 30% (v/v) bleach containing 0.1% (v/v) Triton X-100 and vernalized at 4°C for two days, *Arabidopsis* seeds were germinated on a nylon filter membrane placed on the top of ATS agar plates. The seedlings were grown under continuous fluorescent light (∼80 μmole photon m^-2^ s^-1^) at 22°C for 8 days. For the heat-treated group, these 8-day-old seedlings along with the nlon membrane (mesh opening 20 μm, Cat. #146510, Spectrum Laboratories Inc.,Rancho Dominguez, CA) were then transferred onto fresh ATS [56] media containing 99 atom% K^15^NO_3_ and 98 atom% Ca(^15^NO_3_)_2_ (Cambridge Isotopes Laboratories, Inc., Andover, MA) (^15^N-medium) and then transfer to the 30°C growth chamber. For the control group, seedlings were continuously grown at 22°C after being transferred to the ATS medium with the normal nitrogen source (^14^N-medium).

For both the control and high-temperature groups, crude proteins were extracted at 0, 8, 24, 32, and 48 hours after ^15^N-introduction (time 0 samples was shared by both groups). Prior to transferring seedlings from ^14^N- to ^15^N-medium, ATS liquid medium lacking K^15^NO_3_ or Ca(^15^NO_3_)_2_ was used to rinse the seedlings.

### Proteomic Sample Preparation

For the proteomic analysis of *Arabidopsis* seedlings, hypocotyl and cotyledons (as “shoot” samples) were dissected from root tissues. From root and shoot tissues, soluble and membrane proteins were extracted and enriched by differential centrifugation as described previously by Fan *et al*.[28] in the stable isotope incorporation experiments.. Soluble proteins (150 μg) were precipitated by addition of ice cold acetone to 80% (v/v) followed by overnight incubation at –20°C. The protein precipitate was then pelleted by centrifugation for 15 min at 16,000 g. The air-dried the pellets were dissolved in 8 M urea/8 mM DTT added to a final protein concentration of 8 μg/μL. The proteolysis of soluble protein, membranous protein fractions derived from 1,500 × g (organelle), and 100,000 × g (microsomal) pellets were processed as described previously [28]. The resulting peptides obtained from soluble or membrane protein fractions were purified by C18 solid phase extraction using the C18 Spin column (Pierce Biotechnology, Thermo Scientific, Rockford, IL) and per the manufacturer’s protocol. After purification, peptides were concentrated under vacuum to dryness using a SpeedVac concentrator (Savant) and were re-suspended in 5% (v/v) acetonitrile, 0.1% formic acid prior to ultra-high performance liquid chromatography–high resolution tandem mass spectrometry (UHPLC-HRMS/MS) analysis.

### UHPLC-HRMS/MS Analysis

The tryptic peptides were analyzed by UHPLC-HRMS/MS using a Q Exactive hybrid quadrupole orbitrap mass spectrometer with an Ultimate 3000 UHPLC inlet (Thermo Fisher Scientific, CA) equipped with an ACQUITY UPLC BEH C18 reversed phase column (Waters, 2.1 mm x 100 mm, 1.7 µm particle size). Solvent A (0.1% (v/v) formic acid in H_2_O) and B (0.1% (v/v) formic acid in acetonitrile) were used as mobile phases for gradient separation. The equivalent of 30 μg of soluble protein digest, 10 μg of organellar protein digest or 30 μg of microsomal protein digest were loaded separately onto the column in 5% solvent B for 12 min at a flow rate of 0.3 mL/min, followed by elution by gradient: 2 min from 5% B to 10% B, 60 min to 40% B, 1 min to 85% B and maintained for 10 min. The column was equilibrated for 15 min with 5% B prior to the next run. The MS/MS data were collected in data-dependent acquisition mode similar to Sun *et al.*[57] with minor modifications. Full MS scans (range 350−1800 m/z) were acquired with 70Kresolution. The target value based on predictive automatic gain control (AGC) was 1.0E+06 with 20 ms of maximum injection time. The 12 most intense precursor ions (z ≥ 2) were sequentially fragmented in the HCD collision cell with normalized collision energy of 30%. MS/MS scans were acquired with 35k resolution and the target value was 2.0E+05 with 120 ms of maximum injection time. The ion selection threshold of 1.0E+04 and a 2.0 m/z isolation width in MS/MS was used. The dynamic exclusion time for precursor ion m/z was set to 15 s.

### Protein Identification

All .raw files were converted to mzXML files by msConvert3 tool of ProteoWizard[58] and then converted to mgf format by MGF formatter (v0.1.0). OMSSA (v2.1.9)[59] was used for database searching against the UniProt Arabidopsis thaliana database (accessed on February 2013, 33,311 sequences, http://www.uniprot.org) combined with the contamination list from the cRAP database (common Repository of Adventitious Proteins, accessed on February 2013, 115 sequences, http://www.thegpm.org/crap/) and reversed sequences. The search parameters were: 6 ppm precursor ion mass tolerance, 0.02 m/z product ion mass tolerance, methionine oxidation as variable modification and a maximum missed cleavage of 2. The search results were then analyzed by Scaffold (v3.6.5, Proteome Software Inc., Portland, OR)[60] to validate MS/MS based peptide and protein identifications. The results were filtered with a false discovery rate of less than 0.5% on the peptide level and 1% on the protein level with a minimum of two unique peptides required for identification. Proteins that contained similar peptides and that could not be differentiated based on MS/MS analysis alone were grouped to satisfy the principles of parsimony. All the above activities of data conversion and protein database searching were performed on the Galaxy-P platform (https://galaxyp.msi.umn.edu/),[61–63] and supported by Minnesota Supercomputing Institute of University of Minnesota). The mass spectrometry proteomics data have been deposited to the ProteomeXchange Consortium (http://proteome-central.proteomexchange.org) via the MassIVE partner repository (the dataset identifiers will become available once the manuscript is accepted).

### Calculation of Protein Turnover Rates

The workflow of using the *ProteinTurnover* algorithm is described in the following steps: (1) Data preparation. The Scaffold spectrum report (CSV format) and all MS data (mzXML format) were uploaded for access by the R script; (2) Parameter settings. Parameters such as: stable isotope (^15^N) used for labeling, experimental design (incorporation), peptide ID confidence threshold (80), spectral fitting model (beta-binomial), and nonlinear regression setting(log_2_*k*) were defined; (3) Outputs generated. After finishingthe analysis of a dataset, the results were compiled in a summary html file, which includes model plots (spectral fitting by MLE), EIC plots and regression plots (relative abundance fits) for each individual peptide to be used as needed for manual inspection. The *ProteinTurnover* R script also generates a spreadsheet (.csv) containing peptide turnover information, which includes the peptide amino acid sequences, protein UniProt accession numbers (ID), visual scores, log_2_*k* values and standard errors of log_2_*k*.

For isotope label incorporation experiments, the log_2_ value for each turnover rate constant (log_2_*k*) of each peptide was calculated by performing a non-linear regression of the distribution abundance ratios of unlabeled peptide population against time, assuming a single exponential decay, as previously described in *ProteinTurnover* algorithm [28].

Protein turnover typically exhibits first order kinetics, and the first-order rate constant (*k*) is related to the half-life of the particular peptide by the expression, t_1/2_=(ln(2))/*k*. In this study, the turnover rate was represented by the log_2_*k* values, which are more normally distributed than the untransformed rate constants. After obtaining the turnover results from *ProteinTurnover*, peptides were selected for subsequent inclusion in protein turnover calculations byapplying the following filteringcriteria: (1) the visual score of the spectral fitting(to the beta-binomial model) must be greater than 80; (2) the standard error of the turnover rate must be less than 10; and (3) data must be available for 3 or more time points. The log_2_*k* data of the selected and unique peptides were then averaged to obtain the protein turnover rate.

### Estimating the Difference in log_2_*k* Due to Heat Stress

The selected peptides were analyzed in R to calculate the difference of turnover rate between the control and treated groups. A linear mixed model (LMM) fit with restricted maximum likelihood (using the lme4 package) was applied to estimate the change of protein log_2_*k* between the control and heat-treated group based on peptide log_2_*k* data. The used formula is listed as following:

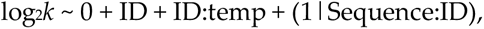

where “ID” represents the protein UniProt accession number, “temp” represents either the control or 30°C group, and “Sequence” represent the peptide amino acid sequence. At the end, only proteins with significant changes in log_2_*k* (p-value less than 0.05) were included in Supplementary Table S-1. Only proteins with more than one computable unique peptide in both control and heat-treated group were selected to generate histograms and box plots (Figure 5, 6, 7).

## 5. Conclusions

This study provides a global look at the dynamics of proteins in plants in response to moderate heat stress. It was conducted at the cellular level with separated soluble and membrane enrichments using ^15^N-stable isotope labeling and the *ProteinTurnover* algorithm for automated data extraction and turnover rate calculation. A total of 571 proteins with a significant change in turnover rates were identified in response to elevated temperature in *Arabidopsis* seedling tissues. Root proteins involved in the redox signaling pathway, stress response, amino acid metabolism, GST metabolism, protein synthesis, protein degradation, and cellular organization appeared to have less change in turnover than shoot proteins. Proteins involved in GST metabolism, photorespiration, protein folding, secondary metabolism, stress response, redox signaling pathway, and beta-glucosidase family proteins exhibited the greatest change in turnover when the temperature was elevated. On the other hand, proteins with the smallest change in turnover were those involved in major carbohydrate metabolism, glycolysis, protein synthesis, and mitochondrial ATP synthesis. This comprehensive study underscores the adaptive mechanisms of plants at the proteomic level under heat stress conditions, potentially guiding future agricultural strategies to enhance crop resilience and productivity in the face of global climate change.

## Author Contributions

Research and investigation, KT.F.; analysis and writing, KT.F and Y.X. All authors have read and agreed to the published version of the manuscript.

## Data Availability Statement

The data presented in this study are available in Supplementary Materials.

## Disclaimer/Publisher’s Note

The statements, opinions and data contained in all publications are solely those of the individual author(s) and contributor(s) and not of MDPI and/or the editor(s). MDPI and/or the editor(s) disclaim responsibility for any injury to people or property resulting from any ideas, methods, instructions or products referred to in the content.

